# Dual inhibition of vacuolar ATPase and TMPRSS2 is required for complete blockade of SARS-CoV-2 entry into cells

**DOI:** 10.1101/2022.03.11.484006

**Authors:** Simoun Icho, Edurne Rujas, Krithika Muthuraman, John Tam, Huazhu Liang, Shelby Harms, Mingmin Liao, Darryl Falzarano, Jean-Philippe Julien, Roman A. Melnyk

**Author notes:** Corresponding author Roman A. Melnyk, Associate Professor of Biochemistry and Senior Scientist, University of Toronto and The Hospital for Sick Children, 686 Bay Street, Toronto, ON M5G 0A4.

## Abstract

An essential step in the infection life cycle of the severe acute respiratory syndrome coronavirus 2 (SARS-CoV-2) is the proteolytic activation of the viral spike (S) protein, which enables membrane fusion and entry into the host cell. Two distinct classes of host proteases have been implicated in the S protein activation step: cell-surface serine proteases, such as the cell-surface transmembrane protease, serine 2 (TMPRSS2), and endosomal cathepsins, leading to entry through either the cell-surface route or the endosomal route, respectively. In cells expressing TMPRSS2, inhibiting endosomal proteases using non-specific cathepsin inhibitors such as E64d or lysosomotropic compounds such as hydroxychloroquine fails to prevent viral entry, suggesting that the endosomal route of entry is unimportant; however, mechanism-based toxicities and poor efficacy of these compounds confound our understanding of the importance of the endosomal route of entry. Here, to identify better pharmacological agents to elucidate the role of the endosomal route of entry, we profiled a panel of molecules identified through a high throughput screen that inhibit endosomal pH and/or maturation through different mechanisms. Among the three distinct classes of inhibitors, we found that inhibiting vacuolar-ATPase using the macrolide bafilomycin A1 was the only agent able to potently block viral entry without associated cellular toxicity. Using both pseudotyped and authentic virus, we showed that bafilomycin A1 inhibits SARS-CoV-2 infection both in the absence and presence of TMPRSS2. Moreover, synergy was observed upon combining bafilomycin A1 with Camostat, a TMPRSS2 inhibitor, in neutralizing SARS-CoV-2 entry into TMPRSS2-expressing cells. Overall, this study highlights the importance of the endosomal route of entry for SARS-CoV-2 and provides a rationale for the generation of successful intervention strategies against this virus that combine inhibitors of both entry pathways.

## Introduction

In December 2019 a novel coronavirus, severe acute respiratory syndrome coronavirus 2 (SARS-CoV-2), emerged causing a global public health crisis [1-4]. The virus quickly spread across the world infecting, as of February 2022, over 430 million people across 220 countries and territories and leading to more than 5.9 million deaths [5]. In early 2021, several vaccines were granted Emergency Use Authorization or approval by the U.S. Food and Drug Administration for their ability to induce an immune response and decrease SARS-CoV-2 infection. However, the emergence of several variants of concern with an altered antigenic profile, a higher infectivity, and a greater likelihood of death threaten the protective efficacy of current vaccines [6-9]. In fact, several studies have already shown a reduction in neutralization potency against these variants by convalescent serum and most monoclonal antibody therapies [10, 11]. Hence, antiviral drugs are urgently needed to provide alternative therapeutic options but also as a safeguard against the emergence of new variants that are poorly managed by current vaccines.

The life cycle of SARS-CoV-2 provides numerous potential avenues for therapeutic intervention. Pfizer’s Paxlovid and Merck’s molnupiravir are two remarkable examples of antivirals that effectively stop viral replication by targeting viral proteins and significantly reducing the risk of hospitalization or death [12, 13]. The SARS-CoV-2 replication cycle starts with the spike (S) protein binding to the cell-surface receptor angiotensin converting enzyme 2 (ACE2) [14, 15]. Then, a cell-surface transmembrane protease, serine 2 (TMPRSS2) cleaves the S protein at the S1/S2 junction and activates the S2 domain [16-19]. The activated S2 domain brings the viral membrane into close proximity to the cell membrane and initiates membrane fusion, which ultimately leads to the release of the viral genome into the host cell [20]. Alternatively, upon ACE2 attachment the virus is endocytosed into endosomes where low-pH-activated proteases such as cathepsin B and cathepsin L activate the S2 domain, which similarly results in viral genome release but in this case through endosomal membrane fusion [21]. Once the viral genome enters the host cytosol it begins propagating [20]. Therefore, targeting any of these steps within the life cycle of SARS-CoV-2 would lead to neutralization of the virus.

Even though TMPRSS2 is expressed on lung and intestinal epithelial cells and, to a lesser extent, in the kidney, heart, adipose, and reproductive tissues, it is not found in other cells susceptible to SARS-CoV-2 infection [22, 23]. Several studies have shown that inhibition of TMPRSS2 activity negatively affects the capacity of SARS-CoV-2 to infect host cells. However, the reported level of attenuation achieved by antivirals targeting TMPRSS2 vary depending on which cell line is used or which variant of SARS-CoV-2 is being tested [16, 24]. More specifically, in the absence of TMPRSS2 expression or in the case of the Omicron variant, antivirals targeting TMPRSS2 are ineffective [16, 25]. Further, Gunst *et al*. demonstrated in a randomised double-blind clinical trial that Camostat, a specific TMPRSS2 inhibitor, did not reduce the time for clinical improvement, progression to intensive care unit admission, or mortality [26]. Therefore, it is unlikely that a monotherapy strategy based on the use of antiviral drugs solely targeting TMPRSS2 will effectively stop viral replication.

Modulating endosomal pH using small molecules represents another attractive strategy that may be employed against SARS-CoV-2. The acidification of endosomes by vacuolar ATPases (V-ATPases) is required for activation of cathepsin proteases, which are ubiquitous across all cells [27, 28]. Although cathepsins represent attractive targets in SARS-CoV-2 infection, potent and/or specific inhibitors of cathepsin B or cathepsin L proteases are not available [17-19]. Furthermore, coronaviruses have been shown to utilize endosomal proteases other than cathepsins for viral activity [29, 30]. Therefore, inhibiting endosomal acidification may represent an optimal strategy to fully neutralize SARS-CoV-2 infection. In fact, recent studies have shown that inhibition of endosomal acidification by using chloroquine or hydroxychloroquine is effective at neutralizing SARS-CoV-2 infection [31]. Yet, the data cannot be replicated in TMPRSS2-expressing cells or translated to the clinic [32-34]. Since both chloroquine and hydroxychloroquine are cationic amphiphilic compounds capable of inducing phospholipidosis, it is unclear which mechanism is operant when neutralizing SARS-CoV-2 infections *in vitro* [35]. By specifically targeting V-ATPases, it may be possible both to neutralize SARS-CoV-2 infection and mitigate off-target effects.

In this study, we set out to investigate the importance of the endosomal pathway for SARS-CoV-2 entry using a collection of small molecule inhibitors of endosomal acidification identified from a high throughput screen. We employed inhibitors using three distinct mechanisms: (1) Lysosomotropic compounds, (2) Proton shuttle compounds, and (3) A direct V-ATPase inhibitor. Our results indicate that widely used non-specific inhibitors of endosomal acidification are toxic at concentrations at which they neutralize the pH, which both confounds interpretation of the mechanism of SARS-CoV-2 inhibition and further portends clinical complications. In contrast, we found bafilomycin A1 to be uniquely potent and safe at inhibiting endosomal acidification and used it to investigate how infection by different variants of SARS-CoV-2 can be blocked by a safe and potent host-targeted mechanism inhibitor. Further, the combination of bafilomycin A1 and Camostat completely and synergistically neutralized SARS-CoV-2 entry into TMPRSS2-expressing cells without confounding cytotoxicity, opening the possibility of a new therapeutic modality that could be both effective and tolerable.

## Results

### Inhibition of endosomal acidification blocks viral entry

To address the importance of endosomal route of entry for SARS-CoV-2 and identify diverse inhibitors of this pathway we developed a robust fluorescence-based high throughput assay endosomal acidification (Supplemental Figure 1A). We screened the MicroSource Spectrum collection of small molecules consisting of 2,360 approved drugs and pharmacologically active molecules with known targets and properties. The top 29 compounds (by percent inhibition) were initially selected for follow-up and tested across a 10-point dose titration for both inhibition of endosomal acidification and compound-mediated toxicity at 24 hours (Supplemental Figure 2A). Among confirmed active compounds that dose dependently inhibited endosomal acidification, in all cases there was overlapping cellular toxicity seen at equivalent doses (Supplemental Figure 2B). To explore this further and determine whether this was a general feature of inhibitors of endosomal acidification, we selected a subset of compounds representing three distinct mechanisms of endosomal deacidification: (1) Lysosomotropic compounds amodiaquine, chloroquine, and quinacrine; (2) Proton shuttle compounds niclosamide and oxyclozanide; and (3) the direct V-ATPase inhibitor bafilomycin A1 (**Figure 1A**). Each of the six compounds were tested for their ability to prevent endosomal acidification, viral entry, and cell viability. Antiviral activity was assessed initially using a pseudotyped virus approach in which the SARS-CoV-2 S protein was pseudotyped onto a human immunodeficiency virus-1 (HIV-1) backbone along with a luciferase reporter gene to measure viral infection (**Figure 1B**) [36]. Inhibition of the pseudoviral particle (PsV) by each compound was tested on HeLa cells expressing ACE2 (HeLa-ACE2). In parallel, each compound was also tested for its ability to inhibit endosomal acidification and its compound-mediated toxicity on HeLa-ACE2 cells over the same dose ranges.

**Figure 1.**
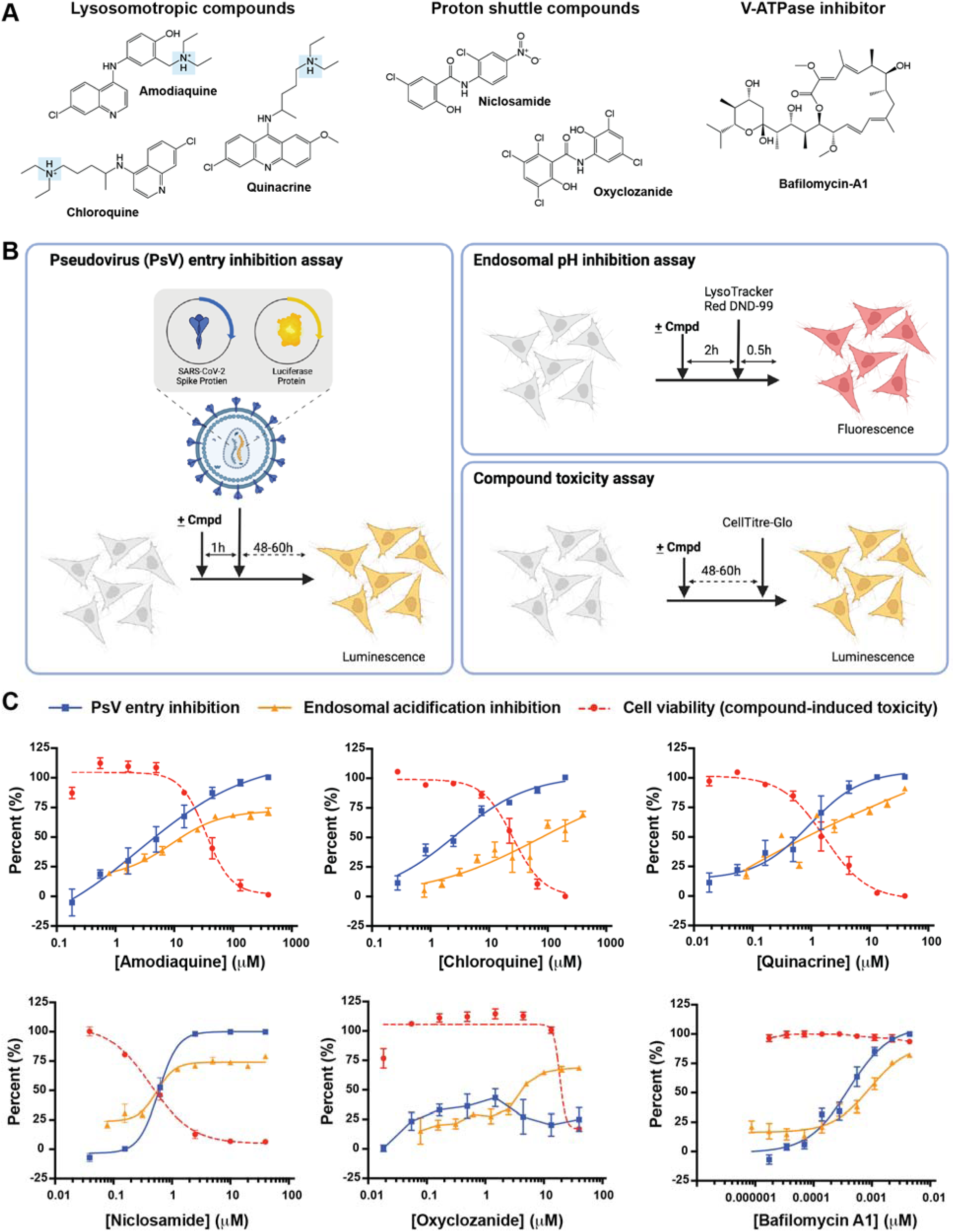
Identification of bafilomycin A1 as a safe and potent inhibitor of SARS-CoV-2 pseudovirus (PsV) entry into mammalian cells. (A) Chemical structure of three groups of endosomal acidification modifiers: lysosomotropic compounds (amodiaquine, chloroquine, and quinacrine), proton shuttles (niclosamide and oxyclozanide), and a specific V-ATPase inhibitor (bafilomycin A1). The highlighted tertiary ammonium cation of the lysosomotropic compounds is a characteristic structure of phospholipidosis-causing compounds. (B) Assays testing pseudoviral entry inhibition (left), endosomal acidification inhibition (top-right), and compound-mediated cell cytotoxicity (bottom-right) of each compound. (C) Comparison of inhibition of PsV entry, inhibition of endosomal acidification, and cell viability by each tested compound. All experiments were conducted in triplicate, at the minimum, (n=3-14).

A reduction in cell infection titers as well as inhibition of endosomal acidification was observed with the three lysosomotropic compounds at a similar concentration range (**Figure 1C**). Specifically, the half-maximal inhibitory concentration (IC_50_) obtained for amodiaquine, chloroquine, and quinacrine was 2.5, 2.6, and 0.9 μM, respectively. However, cell viability significantly decreased as cell infection titers was reduced. The half-cytotoxic concentration (CC_50_) measured for amodiaquine, chloroquine, and quinacrine was 34.7, 25.7, and 1.7 μM, respectively, demonstrating a narrow window in which the compounds can be efficacious against PsVs without causing toxicity. Similarly, incubation of cells with niclosamide, a compound which shuttles protons out of acidified vesicles, at concentrations higher than 1 μM resulted in no measurable infection and efficient endosomal acidification inhibition. However, at those concentrations, cell viability was severely affected (CC_50_ of 0.4 μM). Oxyclozanide, on the other hand, did not alter infection levels nor endosomal acidification at concentrations up to 40 μM but was toxic to cells with a CC_50_ of 18.7 μM (**Figure 1C**). Uniquely, Bafilomycin A1 potently neutralized PsV entry into HeLa-ACE2 cells with an IC_50_ of 0.4 nM and inhibited endosomal acidification with an IC_50_ of 0.9 nM without displaying any associated cell cytotoxicity (**Figure 1C**). These findings highlight fundamental differences among inhibitors of endosomal acidification on compound potency and compound-mediated cellular toxicity.

### Inhibition of either the cell-surface or endosomal entry mechanism attenuates viral entry

In view of its uniquely potent and safe viral inhibition profile, we next sought to use bafilomycin A1 to investigate the relative importance of endosomal activation relative to cell-surface activation [37]. A recent report by Hoffmann *et al*. concluded that endosomal acidification inhibitors, such as chloroquine, are incapable of neutralizing viral entry into cells expressing TMPRSS2 calling into question the importance of the endosomal pathway for viral propagation [32]. For this purpose, we assessed the capacity of bafilomycin A1, alongside the control compound Camostat, to neutralize PsVs on cells expressing ACE2 and TMPRSS2 (Vero-TMPRSS2) and cells expressing ACE2 but lacking TMPRSS2 (HeLa-ACE2) (**Figure 2A**). Consistent with previous reports, Camostat inhibited PsV entry into Vero-TMPRSS2 cells (IC_50_ of 252 nM) but was unable to inhibit PsV entry into HeLa-ACE2 cells lacking TMPRSS2 [32]. By contrast, bafilomycin A1 potently inhibited PsV entry into HeLa-ACE2 cells with an IC_50_ of 0.4 nM and retained its neutralization potency when tested on Vero-TMPRSS2. In fact, bafilomycin A1 was 26-fold more potent than Camostat in stopping viral entry despite the presence of TMPRSS2. These findings are contrary from those reported previously for lysosomotropic molecules, which are suggested to be ineffective in TMPRSS2-positive cells, further highlighting the differences between endosomal inhibitors with distinct mechanisms of action [32].

**Figure 2.**
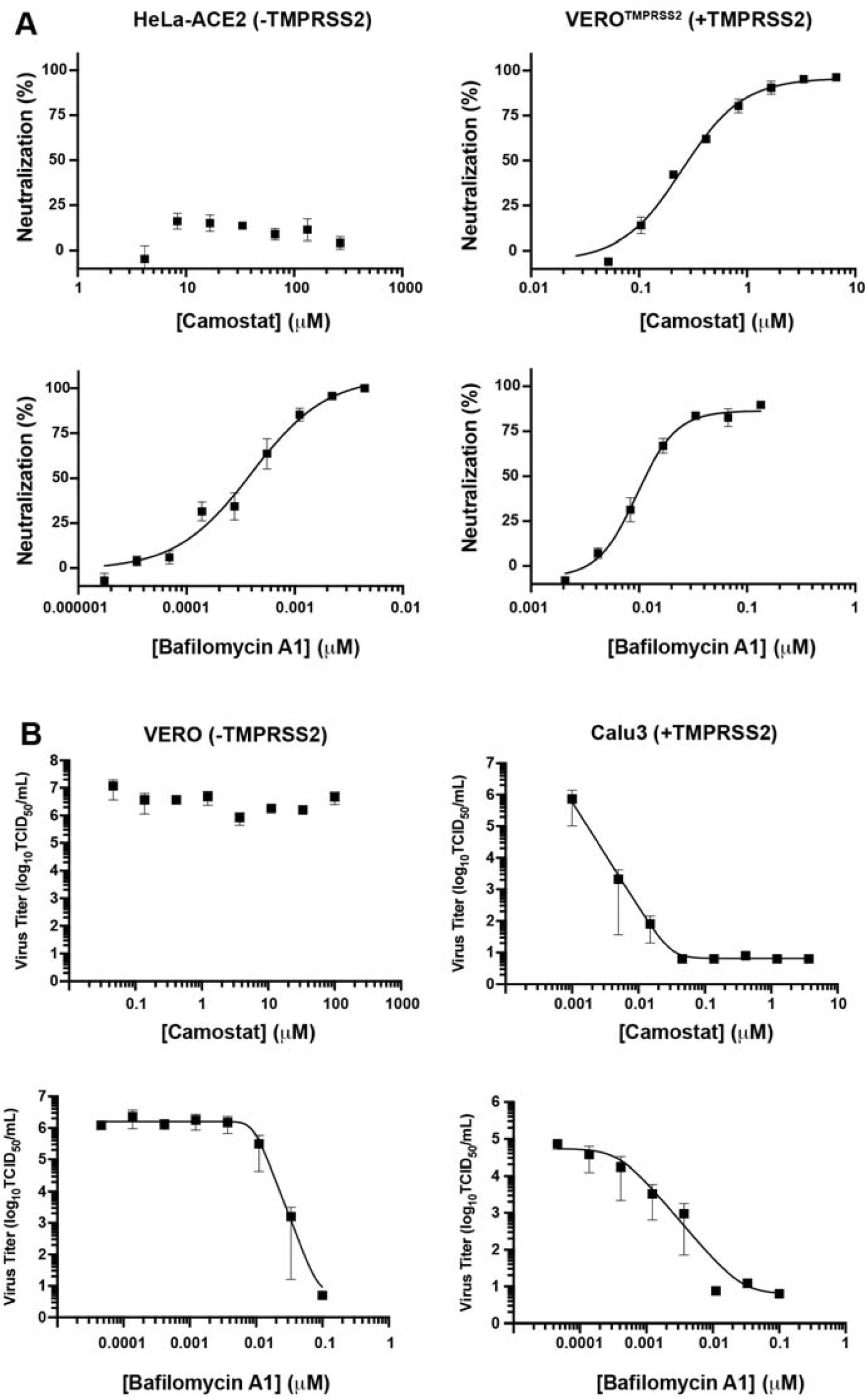
Inhibition of endosomal acidification attenuates SARS-CoV-2 entry irrespective of TMPRSS2 expression. (A) Camostat and bafilomycin A1 were tested against SARS-CoV-2 PsVs using HeLa cells expressing ACE2 but lacking TMPRSS2 or Vero cells expressing both ACE2 and TMPRSS2. All experiments were conducted in at least n=6 up to n=13 (n=6-13). (B) Camostat and bafilomycin A1 were tested against authentic SARS-CoV-2 virus (MOI=0.1) using Vero cells endogenously expressing ACE2 but lacking TMPRSS2 or Calu-3 cells endogenously expressing both ACE2 and TMPRSS2. Data represents three to four independent experiments each done in triplicate using SARS-CoV-2 [SARS-CoV-2/Canada/ON/VIDO-01/2020/Vero’76/p.2 (Seq. available at GISAID – EPI_ISL_425177)] (n=3-4).

To demonstrate that these observations were not cell-dependent, due to TMPRSS2 overexpression, or PsV-specific, we next assayed compound-induced neutralization in Vero cells, which naturally lack TMPRSS2 expression, and Calu-3 cells, which endogenously express TMPRSS2, using authentic SARS-CoV-2 viral particles. Importantly, the above results were reproduced with authentic SARS-CoV-2 viral particles (**Figure 2B**). Consistent with what has been reported previously, Camostat significantly reduced SARS-CoV-2 infection titers in TMPRSS2-expressing Calu-3 cells but not in TMPRSS2-lacking Vero cells [32, 38]. Bafilomycin A1, on the other hand, significantly reduced SARS-CoV-2 infection titers in both Vero and Calu-3 cells. Overall, while inhibition of either entry route effectively inhibits viral infection, inhibition of endosomal acidification via V-ATPase inhibition alone is effective at neutralizing SARS-CoV-2 entry irrespective of TMPRSS2 expression.

### Neutralization of SARS-CoV-2 Variants of Concern by Camostat and bafilomycin A1

Next, we tested whether inhibition of SARS-CoV-2 cell entry by Camostat and bafilomycin A1 is affected by the different variants of the S protein that have emerged, since viral entry efficiency is different across different variants [39]. Specifically, we compared the IC_50_ values of the two compounds against pseudoviruses containing four distinct S proteins: D614G (Wildtype), B.1.1.7 (i.e., Alpha), B.1.351 (i.e., Beta), B.1.1.529 (i.e., Omicron). Camostat inhibited the entry of wildtype, Alpha, and Beta variants into Vero-TMPRSS2 cells; however, was unable to inhibit the entry of the Omicron variant into Vero-TMPRSS2 cells nor, as expected, stop viral infection of any of the variants when tested on HeLa-ACE2 cells lacking TMPRSS2 expression. These findings reinforce the notion that the relative importance of each entry pathway differs among variants and that the Omicron variant relies more heavily on the endosomal route for entry into cells [25]. Consistent with this, bafilomycin A1 inhibited viral entry of all the variants, including the Omicron variant, in both HeLa-ACE2 cells and Vero-TMPRSS2 cells (**Figure 3B**).

**Figure 3.**
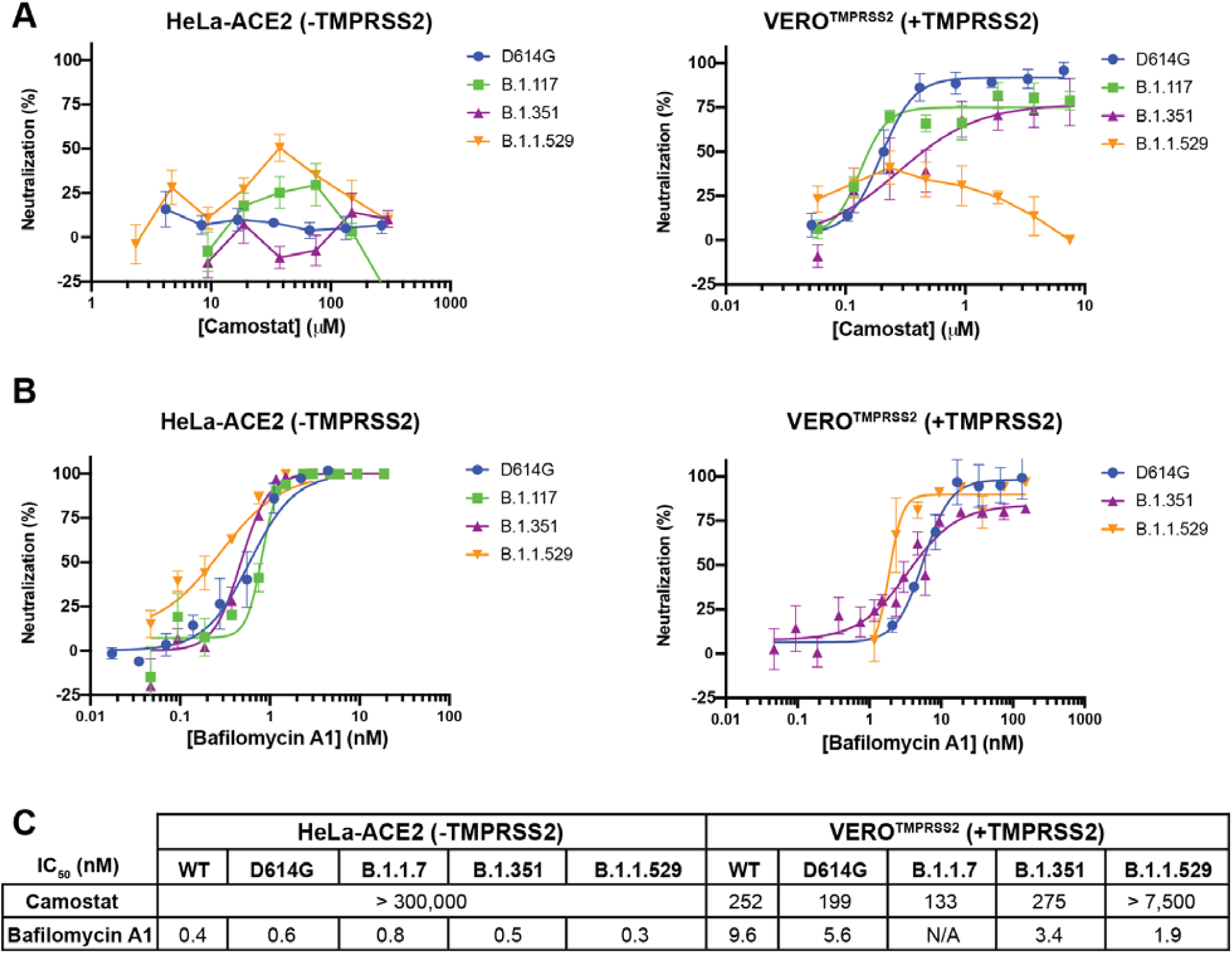
Potency of host-dependent inhibition against different variants of SARS-CoV-2 remains unaffected. Camostat (A) and Bafilomycin A1 (B) were tested against SARS-CoV-2 D641G, B.1.117 (i.e., Alpha), B.1.351 (i.e., Beta) or B.1.1.529 (i.e., Omicron) PsV variants using HeLa cells expressing ACE2 but lacking TMPRSS2 or Vero cells expressing both ACE2 and TMPRSS2. All experiments were conducted in quadruplicates, at the minimum (n=4-5). (C) The half-maximal inhibitory concentration (IC_50_) of Camostat and bafilomycin A1 against several SARS-CoV-2 PsVs variants using HeLa cells expressing ACE2 but lacking TMPRSS2 or Vero cells expressing both ACE2 and TMPRSS2 was calculated using Prism 9 (GraphPad). All experiments were conducted in quadruplicates, at the minimum (n=4-13).

### Camostat and bafilomycin A1 synergistically inhibit SARS-CoV-2 entry

Camostat has been shown to reach maximal blood concentrations of ∼200 nM upon oral dosing [24, 26]. Given the complete lack of protection seen for Camostat on cells lacking TMPRSS2, and only partial protection seen in cells expressing TMPRSS2 (**Figure 2A**), it would be predicted that Camostat would not provide complete blockade of viral entry *in vivo*. Previous studies on the related virus SARS-CoV found that simultaneous inhibition of the endosomal- and cell-surface-entry route was required for complete inhibition of viral entry [40]. Therefore, we hypothesized that a combination of Camostat with bafilomycin A1 would inhibit both entry pathways and thus lead to the complete inhibition of SARS-CoV-2 entry. To this end, we tested two doses of bafilomycin A1 that inhibit viral entry (*viz*. 8.3 and 16.7 nM) in combination with a range of Camostat doses covering a full dose titration on Vero-TMPRSS2 cells and measured PsV entry (**Figure 4A**). Across all doses of Camostat, both doses of bafilomycin A1 significantly increased the inhibition of PsV entry into TMPRSS2-expressing cells as compared to Camostat alone.

**Figure 4.**
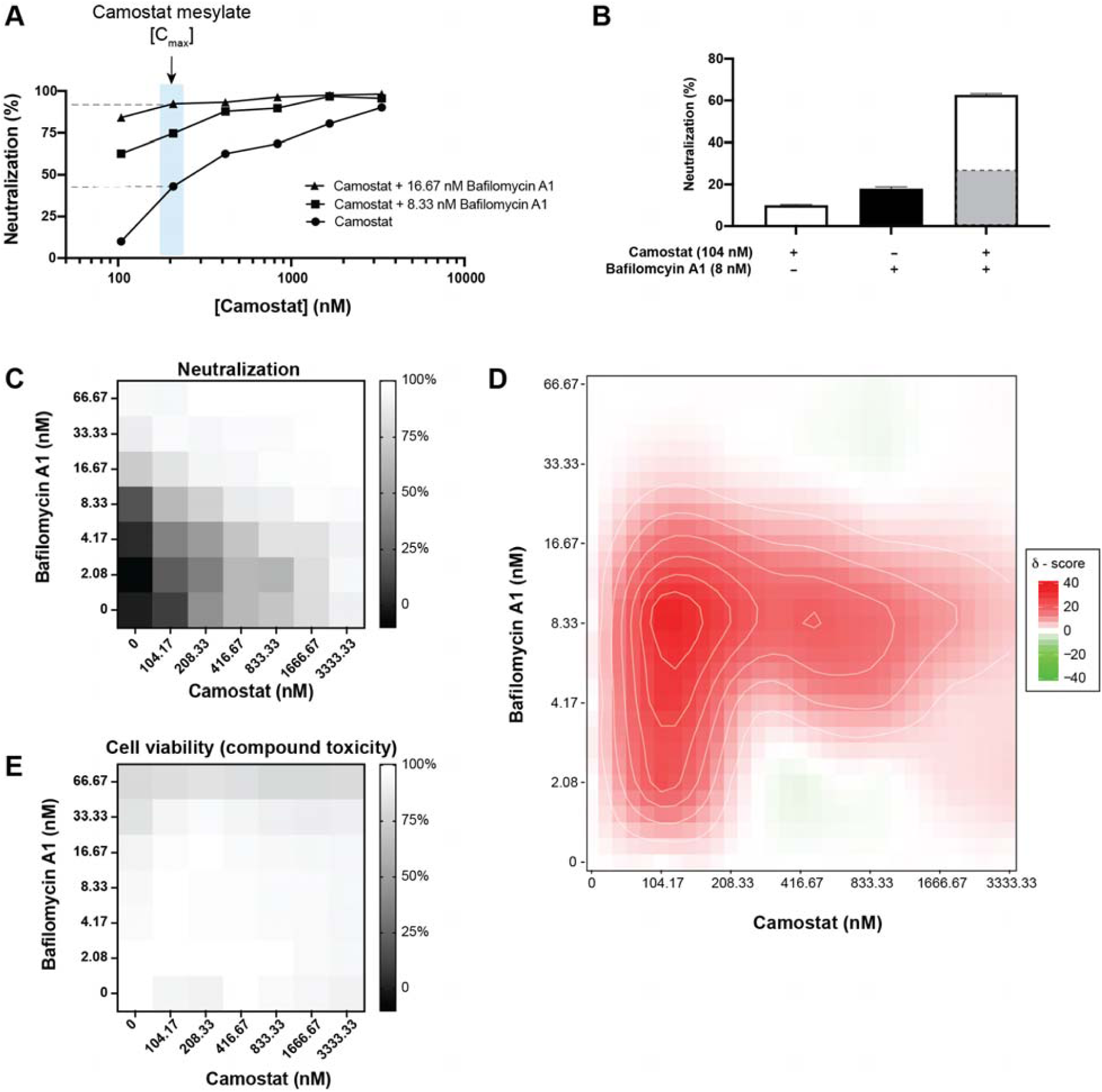
Host-dependent dual-entry inhibition safely and synergistically inhibits SARS-CoV-2 pseudovirus entry into mammalian cells. (A) A dose-dependent inhibition of SARS-CoV-2 PsV entry into Vero cells expressing both ACE2 and TMPRSS2 was measured using Camostat alone or Camostat in addition to either 8 nM or 17 nM bafilomycin A1. The blue box highlights the maximal blood concentration Camostat was found to achieve in patients [26]. All experiments were conducted in duplicate (n=2). (B) 104 nM Camostat and 8 nM bafilomycin A1 were tested individually or in combination against SARS-CoV-2 PsVs using Vero cells expressing both ACE2 and TMPRSS2. The highlighted gray bar in the combination experiment highlights the theoretical neutralization if the two compounds acted additively. All experiments were conducted in duplicate (n=2). (C) A checkerboard assay using Camostat and bafilomycin A1 was conducted against SARS-CoV-2 PsV using Vero cells expressing both ACE2 and TMPRSS2. The experiments were conducted in duplicate (n=2). (D) A Zero Interaction Potency (ZIP) score of the checkboard assay using Camostat and bafilomycin A1 from (C) was calculated using SynergyFinder 2.0. (E) A checkerboard assay using Camostat and bafilomycin A1 at the same concentrations as (C) on Vero cells expressing both ACE2 and TMPRSS2 was tested for compound-mediated toxicity. The experiment was conducted in triplicate (n=3).

The addition of 8.33 nM bafilomycin A1 to 104 nM of Camostat drastically improved neutralization of SARS-CoV-2 beyond what would be expected if the two compounds inhibited additively (**Figure 4B**). In view of these results, we explored whether the simultaneous inhibition of the two entry mechanisms of SARS-CoV-2 using Camostat and bafilomycin A1 could synergistically inhibit viral entry. To assess this and following a checkerboard assay format, we employed the Zero Interaction Potency (ZIP) model which combines the Bliss independence model and the Loewe additivity model [41]. The Bliss independence model assumes each compound elicits its effect independently, while the Loewe additivity model assumes both compounds are the same and thus the activity must be double when two compounds are combined at equal ratios. Hence, combined, the ZIP model identifies potential synergy between two compounds by comparing the change in potency of each compound with different combinations. A ZIP synergy score >10 has been proposed as a threshold indicative of synergistic inhibition by two compounds [41]. Using this analysis, we observed that a combination of 104 nM of Camostat and either 2.08 nM, 4.16 nM, 8.33 nM, or 16.67 nM of bafilomycin A1 had a synergy score of 23.2, 28.5, 34.7, and 15.6, respectively. At 8.33 nM bafilomycin A1, the synergy score was >16 as the concentration of Camostat increased from 104 nM to 833 nM indicating that this combination is highly synergistic at inhibiting PsV entry into Vero-TMPRSS2 cells (**Figure 4D**). In addition, the data shows that neither Camostat nor bafilomycin A1 alone fully inhibited PsV entry even at the highest concentrations tested (3333 nM and 66.7 nM, respectively) but, when combined, the two compounds provided complete inhibition (**Figure 4C**). Importantly, none of the combinations had a toxic effect on Vero-TMPRSS2 cells (**Figure 4E**). Taken together, by simultaneously blocking without cytotoxicity both entry mechanisms using Camostat and bafilomycin A1, SARS-CoV-2 is completely inhibited in a synergistic manner from infecting mammalian cells.

## Discussion

The prospect of novel SARS-CoV-2 variants of concern emerging that are more transmissible and/or are resistant to the antibodies raised by natural infection, through vaccination or developed as therapeutics underscores the need for developing antivirals acting through distinct mechanisms [6-9]. Promising phase III clinical trial data from Pfizer’s Paxlovid and Merck’s Molnupiravir have demonstrated the potential of small molecule therapeutics against SARS-CoV-2 [12, 13]; however, it remains to be seen whether resistance against these virus targeted compounds will emerge [42]. An alternative strategy to block the virus lifecycle is to target the host factors and host mechanisms that are required for SARS-CoV-2 infection. This strategy has the added benefit of being resilient to SARS-CoV-2 mutations since the likelihood of drug resistance against a host-dependent mechanism is negligible without substantial changes to the S protein structure and function [43]. Here, we focused on the major categories of host proteases implicated in the two distinct viral entry routes used by SARS-CoV-2: (1) The cell-surface route (mediated by TMPRSS2 found in some but not all cells), and, (2) the endosomal route (primarily mediated by cathepsin B and cathepsin L, found in all cells) [28]. Because endosomal cathepsins require the low pH of acidified endosomal compartments for activation, it has been recognized that agents that raise the pH of endosomes can indirectly inhibit both cathepsin B and cathepsin L [27].

In the present study, we screened a panel of prototypic inhibitors of endosomal acidification that act through three distinct mechanisms to identify the most efficacious means to inhibit viral entry across multiple cellular contexts. Given recent reports of confounding toxicities for a subset of molecules that inhibit endosomal pH as part of their mechanism [35], together with a lack of clinical efficacy seen for molecules like hydroxychloroquine (a lysosomotropic molecule) [33, 34], we measured in parallel the extent to which each of the molecules tested induced cellular toxicity. We found that both lysosomotropic compounds (amodiaquine, chloroquine, and quinacrine) and a compound capable of shuttling protons out of the endosome (niclosamide) were cytotoxic at doses perceived to be effective at neutralizing viral entry. By contrast, we found that inhibition of endosomal acidification through a specific V-ATPase inhibitor (bafilomycin A1) was safe and effective at neutralizing SARS-CoV-2 entry into cells. The larger safety window of bafilomycin A1 minimally suggests that not all modes of inhibiting endosomal acidification are equal but more importantly demonstrates that the concomitant cytotoxicity of all other compounds confounds their further use to understand how SARS-CoV-2 is neutralized. Importantly, in support of this mechanism being important for viral entry, Daniloski *et al*., and others, used CRISPR (clustered regulatory interspaced short palindromic repeats) knockout screens to demonstrate that the disruption of the V-ATPase or endosomal acidification pathway results in protection against SARS-CoV-2 infection of mammalian cells [44, 45].

In contrast to Camostat, which only protects cells expressing TMPRSS2, we found that bafilomycin A1 was active in all cell lines tested irrespective of TMPRSS2 expression. These results emphasize the importance of the endosomal pathway in the cell entry of SARS-CoV-2, a mechanism previously suggested not to be important in the presence of TMPRSS2 [32]. These contradictory findings might be due, at least in part, to the cytotoxicity associated with compounds such as chloroquine or hydroxychloroquine that have been previously used to elucidate the importance of endosomal acidification for the entry of SARS-CoV-2. Indeed, their compound-mediated toxicity and different modes of inhibiting SARS-CoV-2 propagation makes it difficult to delineate the importance of endosomal acidification on viral entry [46-48]. Further, recent work by Tummino *et al*. demonstrated that cationic amphiphilic compounds, such as chloroquine, induce phospholipidosis at efficacious doses underscoring the need for careful assessment of off-target toxicity of compounds with multiple modes of action [35]. Taken together, safely and specifically inhibiting endosomal acidification using bafilomycin A1 blocks viral entry into mammalian cells irrespective of TMPRSS2 expression.

A recent report has demonstrated a decreased reliance on TMPRSS2 and a concurrent increased dependence on the endosomal route of entry by the Omicron variant, as compared to other variants such as Delta [25]. This shift highlights both the importance of the endosomal route of entry and the need for an antiviral drug strategy that will remain effective as SARS-CoV-2 continues to mutate and evolve. In line with these findings, Camostat was able to inhibit most of the tested variants with a similar efficacy as compared to the wildtype virus using Vero-TMPRSS2 cells but was unable to inhibit viral entry of the Omicron variant. More importantly, we found that bafilomycin A1 inhibited all tested variants with a similar efficacy as compared to the wildtype pseudovirus, and did so irrespective of TMPRSS2 expression. Therefore, targeting endosomal acidification specifically appears to be the strategy by which multiple variants of concern are potently inhibited.

Here, we show that a combination of Camostat and bafilomycin A1 fully inhibit viral entry in TMPRSS2-expresing cells in a synergistic manner and without cytotoxicity. In a recent clinical trial, Camostat was shown to be tolerable at oral doses which can achieve a blood concentration of ∼200 nM [26]. Unfortunately, Camostat on its own was shown to be ineffective at protecting against SARS-CoV-2 infections and bafilomycin A1 has yet to be progressed to a clinical trial as a single compound treatment. Our study identifies a new modality that should be explored in future experiments where a cell-surface inhibitor and endosomal route inhibitor are simultaneously used to protect against SARS-CoV-2 infections. Supporting this idea, Yuan *et al*. demonstrate the power of using two inhibitors targeting different mechanisms of the SARS-CoV-2 life cycle to synergistically inhibit SARS-CoV-2 infection *in vivo* [49]. Importantly, inhibition of the endosomal route of entry, using bafilomycin A1, equally neutralized all variants tested. Our results suggest that the use of such inhibitors targeted at host-dependent mechanisms may be resilient against emerging variants of concern, and capable of broadly blocking viral entry into mammalian cells

## Materials And Methods

### High Throughput Screen of Endosomal Acidification

Inhibition of endosomal acidification was measured using LysoTracker Red DND-99 (ThermoFisher). Vero cells (ATCC) were seeded in 96-well clear CellBind plates (Sigma Aldrich) 24 hours prior to the experiment at a density of 40,000 cells/well in a 100 μL volume of complete DMEM supplemented with 10% inactive FBS and 1% of penicillin/streptomycin. On the day of the experiment, the media was replaced with 100 μL of serum free DMEM. A volume of 0.4 μL of each compound from the Spectrum Collection library (MicroSource), consisting of 2,560 individual compounds formatted as 10 mM solutions in DMSO, was incubate with the cells at 37°C for 2 hours. Following this, 0.1 μM final concentration of LysoTracker Red DND-99 (ThermoFisher) was added to each well and incubated for 30 minutes at 37°C. The cell media was then replaced with 100 μL of FluoroBrite DMEM (ThermoFisher). Fluorescence at excitation/emission 574/594 nm was measured using an Envision plate reader (Perkin Elmer). Data was plotted using Prism 9 (GraphPad).

### Pseudovirus production and Neutralization assay

HIV-1-derived viral particles were pseudotyped with full-length SARS-CoV-2 spike (S) protein as described previously [50]. Briefly, plasmids expressing a lentiviral backbone encoding the luciferase reporter gene (BEI NR52516), the HIV structural and regulatory proteins Tat (BEI NR52518), Gag-pol (BEI NR52517) and Rev (BEI NR52519), and the full-length SARS-CoV-2 S protein were co-transfected into human kidney HEK293T cells (ATCC, CRL-3216) using the BioT transfection reagent (Bioland Scientific) following the manufacturer’s instructions. In order to test the effect of the S protein mutation D614G that renders a more infectious virus [8], the plasmid encoding the S protein containing the D614G mutation (kindly provided by D.R. Burton; The Scripps Research Institute) was used instead of the plasmid encoding the wild-type SARS-CoV-2 S protein. Similarly, the wild-type S plasmid was substituted with plasmids codifying for the B.1.117 or B.1.351 SARS-CoV-2 S proteins (kindly provided by David Ho, Columbia University) or B.1.1.529 (synthesized and cloned by GeneArt, LifeTechnologies into pcDNA3.4 expression vector) to generate the corresponding SARS-CoV-2 pseudoviral particle (PsV) variants. Transfected cells were incubated at 37°C and after 24 hours, 5 mM sodium butyrate was added to the media. Cells were incubated for an additional 24-30 hours at 30°C, after which PsVs were harvested, passed through 0.45 μm pore sterile filters, and finally concentrated using a 100K Amicon Ultra 2.0 Centrifugal Filter Units (Merck Millipore Amicon).

Neutralization was determined in a single-cycle neutralization assay using HeLa cells expressing ACE2 (HeLa-ACE2), kindly provided by D.R. Burton; The Scripps Research Institute, and Vero E6 cells constitutively expressing the transmembrane protease, serine 2 (Vero-TMPRSS2), obtained from the Centre For AIDS Reagents (National Institute for Biological Standards and Control) [51, 52]. Cells were seeded in 96-well clear CellBind plates (Sigma Aldrich) 24 hours prior to the experiment at a density of 10,000 cells/well in a 100 μL volume. On the day of the experiment, the sample compounds were serially diluted in complete DMEM media (cDMEM^2%^) that contained 2% inactive fetal bovine serum (FBS) and 50 μg/mL of gentamicin. Media from the cell culture was replaced with 100 μL of fresh cDMEM^2%^ media and 50 μL of the previously diluted sample compounds were added to each well and incubated for 1 hour at 37°C. After incubation, 50 μL of PsVs was added to each well and incubated for 48-60 hours in the presence of 10 μg/mL of polybrene (Sigma Aldrich). Unless stated otherwise, PsV baring the SARS-CoV-2 S protein (BEI NR52310) was used. PsV entry level was calculated based on luminescence in relative light units (RLUs). For that, 130 μL of supernatant was aspirated from each well to leave approximately 50 μL of media. 50 μL Britelite plus reagent (PerkinElmer) was added to each condition and incubated for 2 minutes at room temperature. 100 μL volume was transferred to a 96-well white plate (Sigma-Aldrich) and luminescence was read using a Synergy Neo2 Multi-Mode Assay Microplate Reader (Biotek Instruments). Data was plotted and half-maximal inhibitory concentration (IC_50_) values were calculated using Prism 9 (GraphPad).

In order to confirm that the reduced luminescence was not related to cell toxicity, HeLa-ACE2 and Vero-TMPRSS2 cell viability upon incubation with serial dilutions of the sample compounds were tested in parallel to the neutralization assay. Hence, following the above-mentioned protocol, 10,000 cells/well of pre-seeded cells were co-cultured with the same serial dilutions of the sample compounds at 37°C for 48-60 hours. Cell viability was monitored by adding 50 μL of CellTiter-Glo reagent (Promega) to 200 μL of media containing cells. After a 10-minute incubation, 100 μL volume was transferred to a 96-well white plate (Sigma-Aldrich) to measure luminescence using a Synergy Neo2 Multi-Mode Assay Microplate Reader (Biotek Instruments). Data was plotted and half-cytotoxic concentration (CC_50_) values were calculated using Prims 9 (GraphPad).

Inhibition of endosomal acidification was measured using LysoTracker Red DND-99 (ThermoFisher). HeLa-ACE2 and Vero-TMPRSS2 cells were prepared following the above-mentioned protocol, where 10,000 cells/well of pre-seeded cells were co-cultured with the same serial dilutions of the sample compounds at 37°C for 2 hours. Following this, 0.1 μM final concentration of LysoTracker Red DND-99 (ThermoFisher) was added to each well and incubated for 30 minutes at 37°C. The cell media was then replaced with 100 μL of FluoroBrite DMEM (ThermoFisher). Fluorescence was measured using a Synergy Neo2 Multi-Mode Assay Microplate Reader (Biotek Instruments) while employing a Texas Red filter. Data was plotted and IC_50_ values were calculated using Prism 9 (GraphPad).

### Authentic SARS-CoV-2 Neutralization and Titer Assay

Authentic SARS-CoV-2 experiments were conducted using Vero’76 (ATCC, CRL-1587) or Calu-3 (ATCC, HTB-55) cells. Cells were seeded and grown in complete DMEM media (cDMEM) that contained 10% inactive FBS and 1% of penicillin/streptomycin overnight at 37°C to approximately 90% confluence in 96-well plates. SARS-CoV-2 [SARS-CoV-2/Canada/ON/VIDO-01/2020/Vero’76/p.2 (Seq. available at GISAID – EPI_ISL_425177)] virus was diluted in complete DMEM supplemented with 2% inactive FBS and 1% of penicillin/streptomycin to obtain a multiplicity of infection (MOI) of 0.1 (approximately 2000 TCID_50_/well). Two test compounds, Camostat and bafilomycin A1, were prepared to a 3-fold dilution in DMEM supplemented with 2% inactive FBS. Cells were incubated with the test compounds for 1 hour prior to infection at which point 50 μL of viral inoculum was added to each well. After 1 hour of incubation with the viral inoculum, the inoculum and compound mixture were removed and fresh compounds were added to the plate. The plates were incubated for 48 hours at 37°C. At 24 hours, plates were assessed for contamination. Media alone and cell-alone control wells (without viral infection and without compound treatment) were used as controls for virus replication.

The plate of infected cells that had been exposed to the test compounds was examined for cytopathic effect (CPE) and cytotoxicity (if noticeable) under a microscope at 48 hours. At 48 hours, 100 μL of supernatant from each well was transferred into 96-well rounded-bottom plates. Viral titration by the median tissue culture infectious dose (TCID_50_) assay of the supernatant was carried out by a serial 7-fold dilution. These dilutions were used to infect pre-seeded cells, as described for the initial infection. Cells were observed for CPE at 1, 3 and 5 days after the infection. The Spearman-Karber algorithm were used to calculate TCID_50_ titers which were used to calculate IC_50_ values for each compound using Prims 9 (GraphPad).

### Pseudovirus Neutralization Synergy Assay

Synergistic neutralization by a combination of two compounds was defined as an increase in the inhibition of PsV entry upon the combination of two different compounds in comparison to the sum of the expected inhibition of both molecules or double the concentration of the best molecule in the *in vitro* assay [41]. Hence, different ratios of the two compounds were mixed and serial dilutions were prepared in a total volume of 50 μL for each condition using a checkerboard assay. The assay was performed using Vero-TMPRSS2 cells following the above protocol. The level of inhibition of each condition was calculated and compared to the expected values using the Zero Interaction Potency (ZIP) synergy score model [41]. Synergy scores were assigned for each condition and values above 10 were interpreted as a synergistic effect.

In order to confirm that a combination of two compounds did not also lead to increased cell toxicity Vero-TMPRSS2 cell viability was measured using CellTiter-Glo reagent (Promega). Vero-TMPRSS2 cells were seeded overnight at 10,000 cells/well in 96-well clear CellBind plates (Sigma Aldrich). Media was exchanged with 100 μL of serum-free DMEM containing 1% penicillin-streptomycin. An Agilent Bravo liquid handler was used to deliver 0.27 μL of bafilomycin A1 then 0.27 μL of Camostat to the cell plates. The cell plates were incubated at 37°C for 48-60 hours before 50 μL of CellTiter-Glo reagent (Promega) was added. Cell plates were gently mixed at room temperature for 10 minutes then 100 μL of the mixture was transferred to 96-well white plates to measure luminescence using a SpectraMax M5 (Molecular Devices). Data was plotted using Prism 9 (GraphPad).

## ACKNOWLEDGEMENTS

This research was funded (SI and RAM) from Fast Grants, part of the Emergent Ventures Program at the Mercatus Centre at George Mason University, with support from Thistledown Foundation. This research was supported by the European Union’s Horizon 2020 research and innovation program under Marie Sklodowska-Curie grant 790012 (ER). This work was further supported by the CIFAR Azrieli Global Scholar program (JPJ), the Ontario Early Researcher Award program (JPJ) and the Canada Research Chair program (JPJ). The Synergy Neo2 Multi-Mode Assay Microplate Reader instrument was accessed at the Structural and Biophysical Core Facility, The Hospital for Sick Children, supported by the Canada Foundation for Innovation and Ontario Research Fund.

## Notes

### Competing Interest Statement

The authors have declared no competing interest.

