## Supplemental Files for "Dual inhibition of vacuolar ATPase and TMPRSS2 is required for complete blockade of SARS-CoV-2 entry into cells"

Supplemental Figure 1

A)

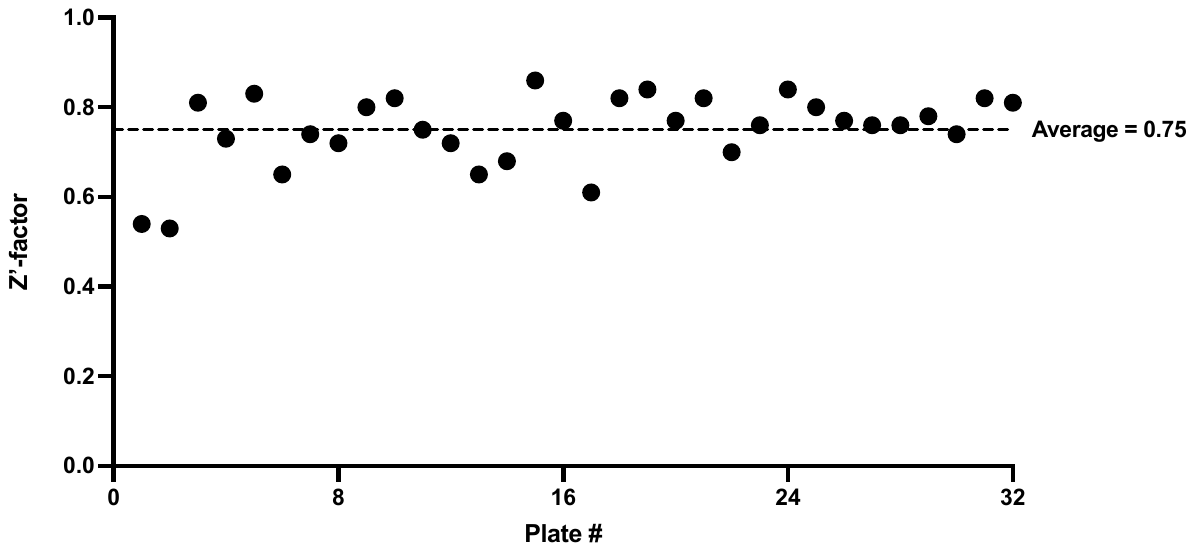

B)

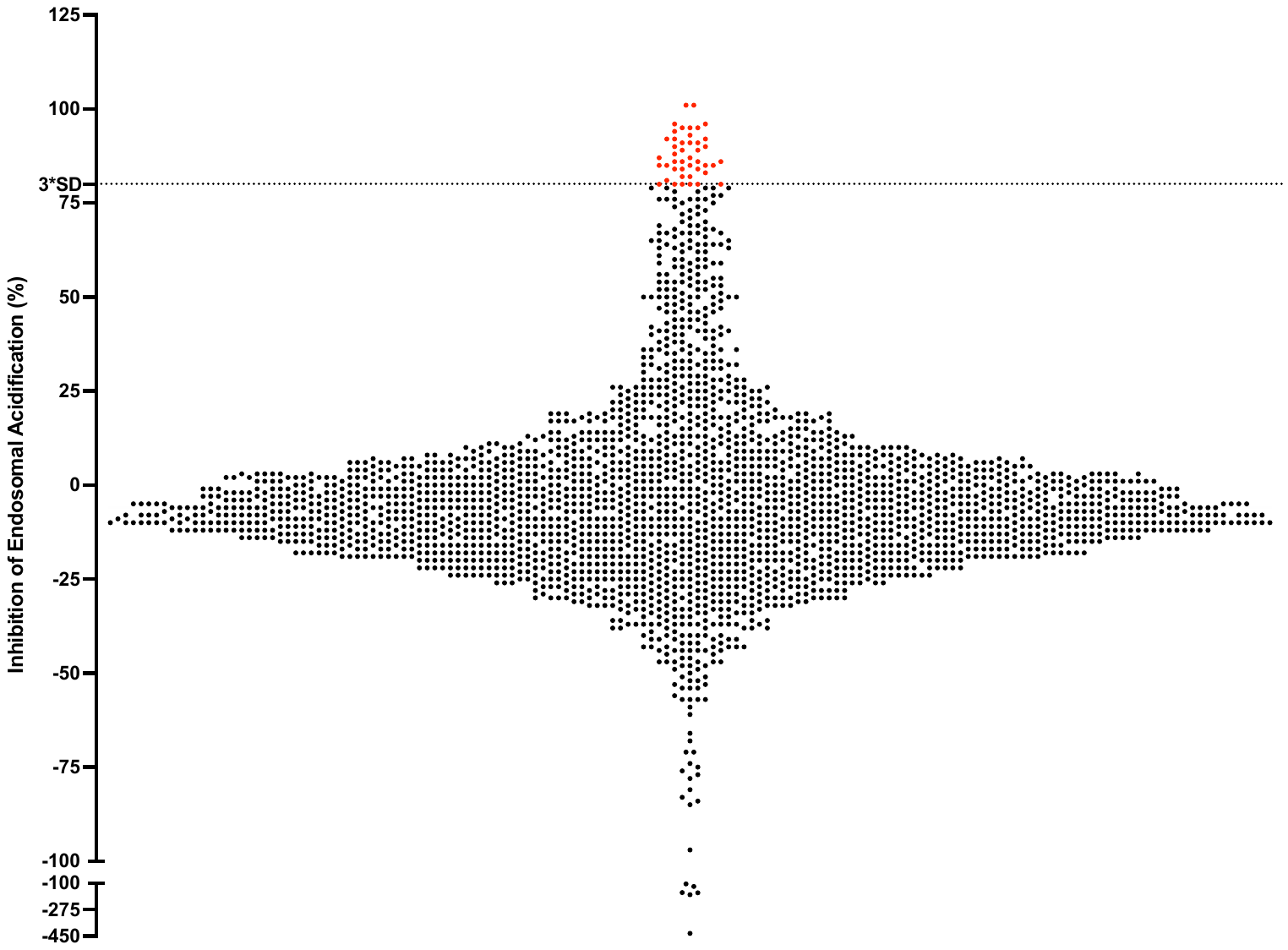

Supplemental Figure 2

A)

| **Compound Name** | **Structure** |
| --- | --- |
| Agaric acid | 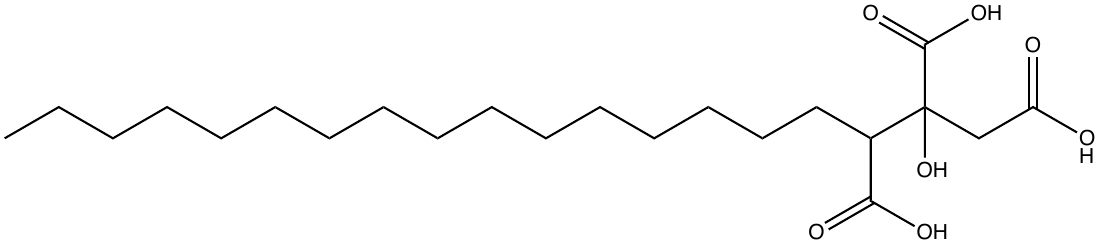 |
| Alexidine | 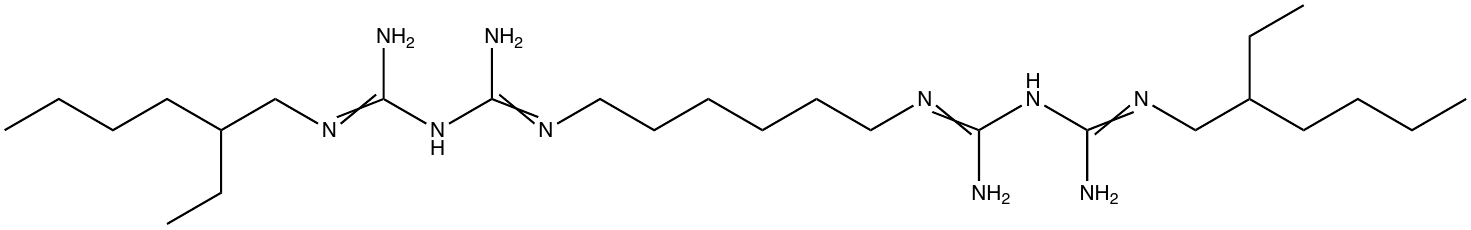 |
| Amsacrine | 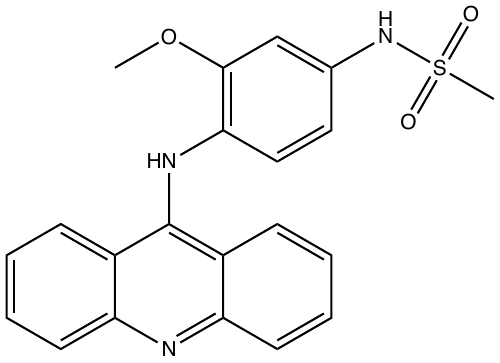 |
| Bazedoxifene | 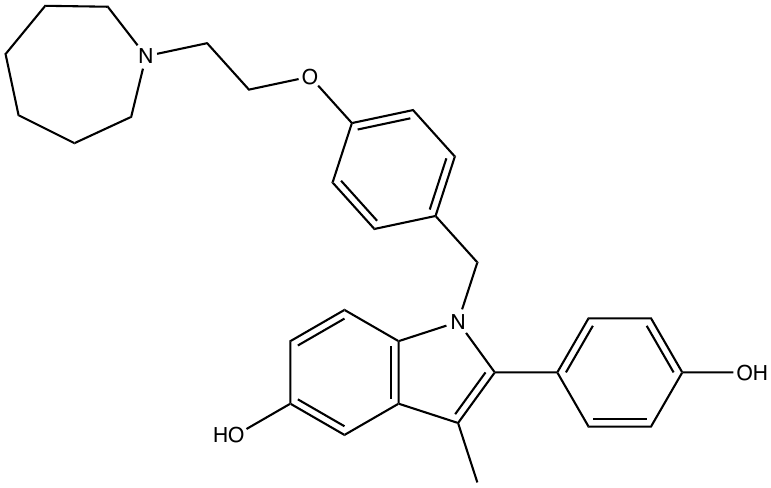 |
| Benzalkonium chloride | 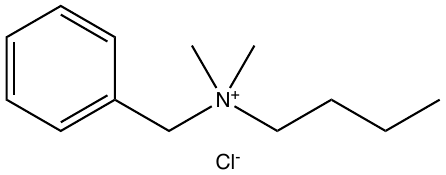 |
| Benzethonium chloride | 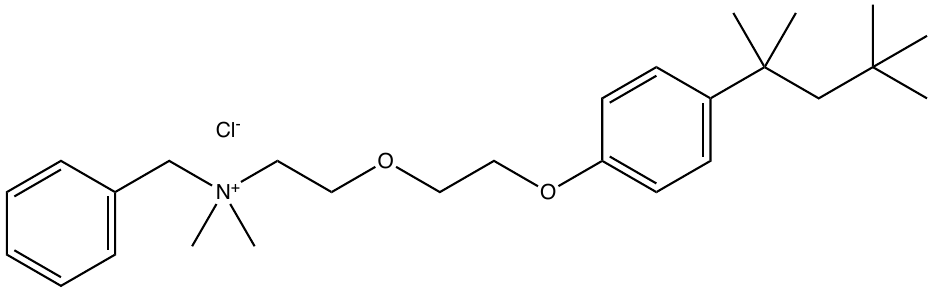 |
| Celastrol | 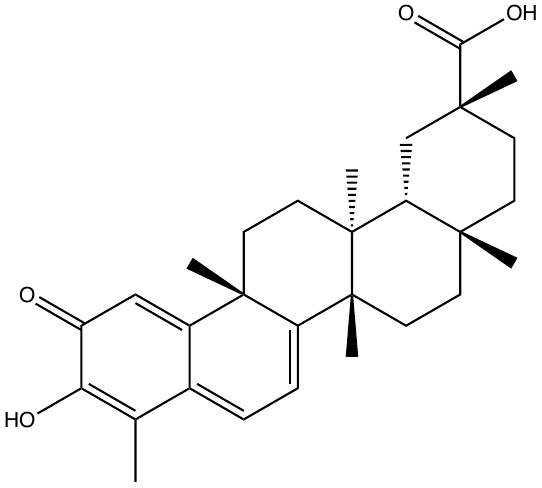 |
| Cetalkonium chloride | 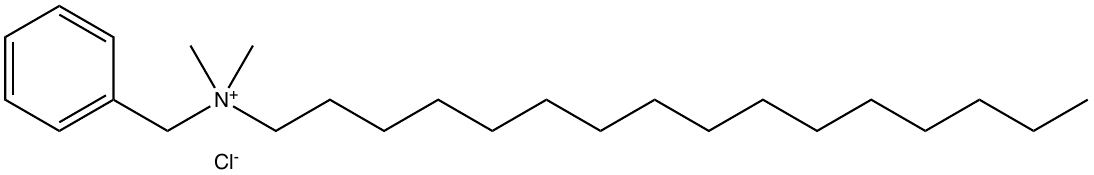 |
| Cinacalcet hydrochloride | 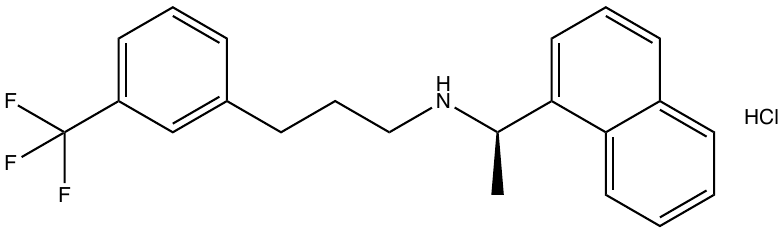 |
| Clofazimine | 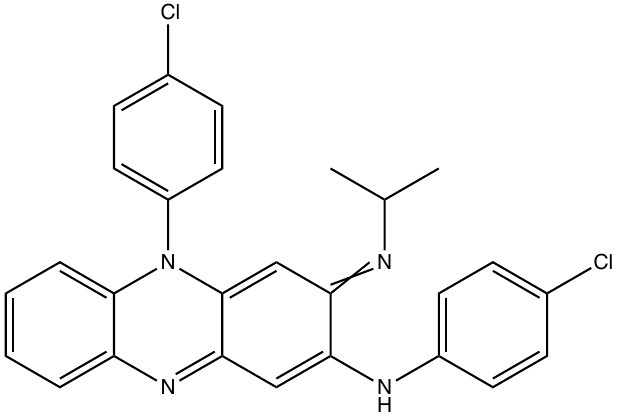 |

| **Compound Name** | **Structure** |
| --- | --- |
| Clomipramine | 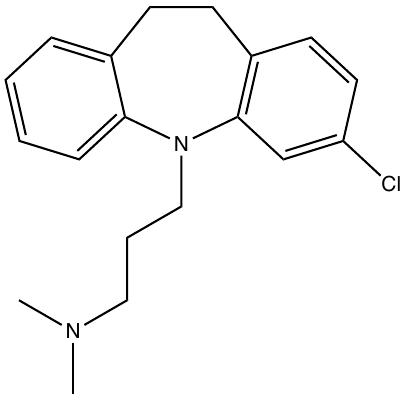 |
| Dapoxetine hydrochloride | 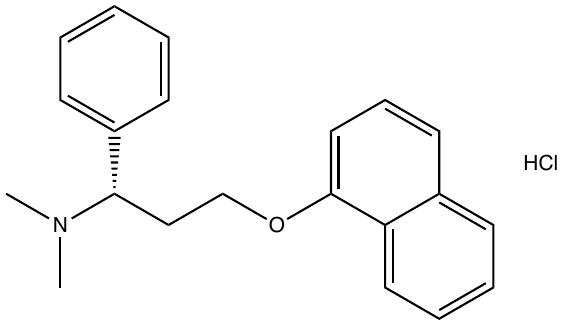 |
| Dronedarone | 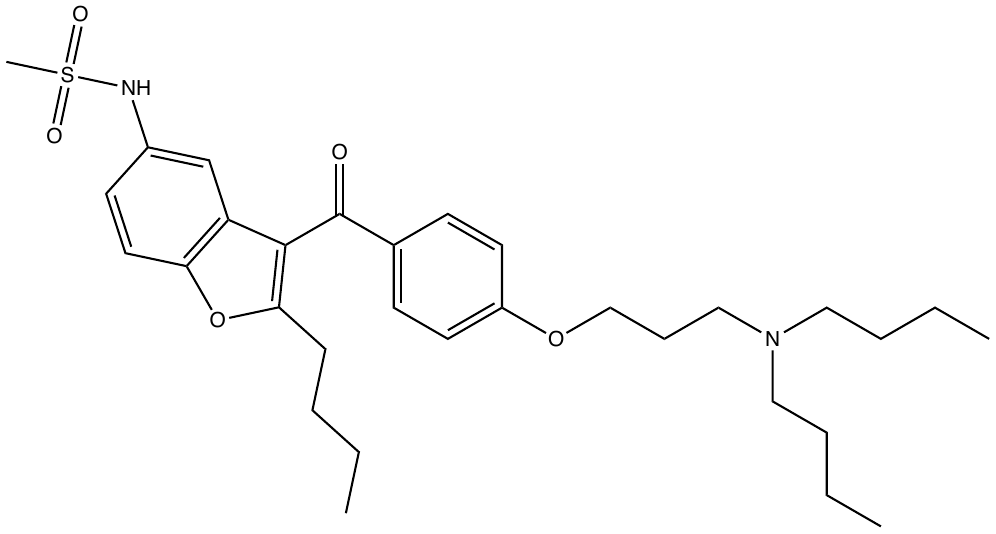 |
| Gambogic acid | 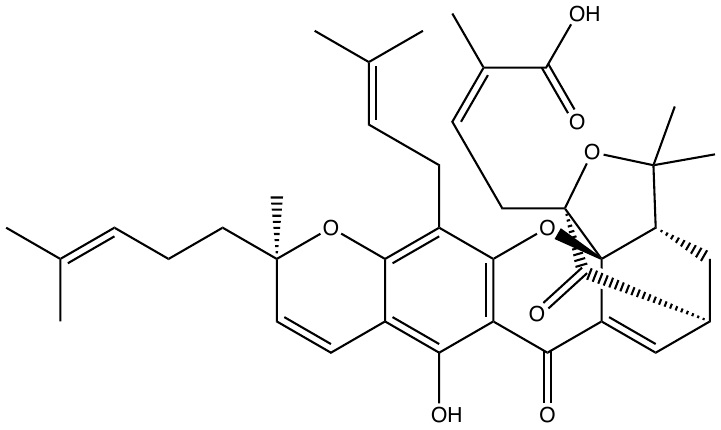 |
| Gossypol | 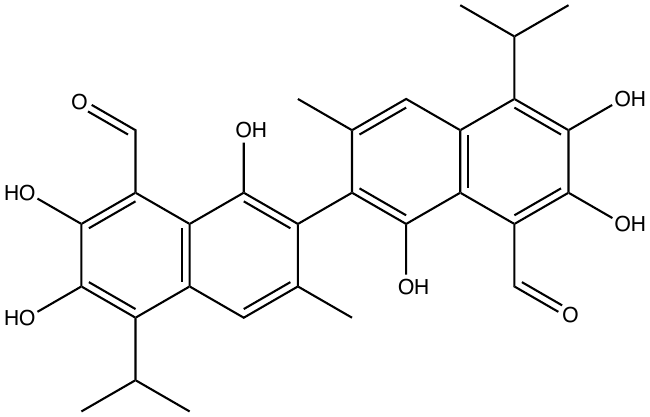 |
| Indapamide | 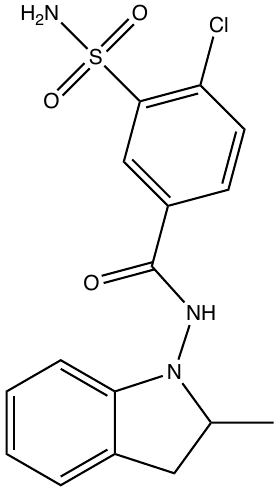 |
| Methylbenzethonium chloride | 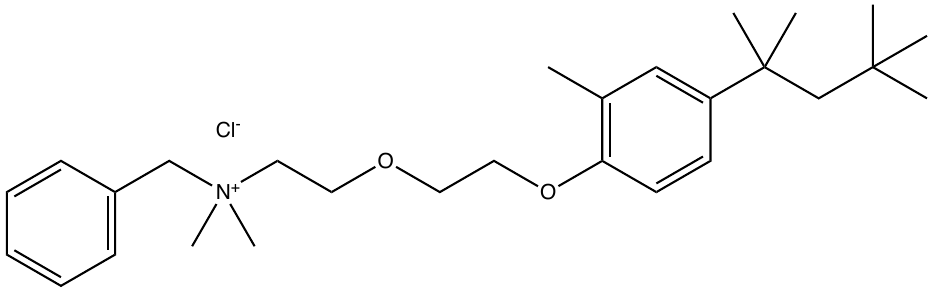 |
| Methylene Blue | 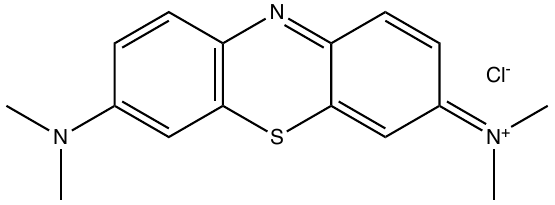 |
| Miconazole | 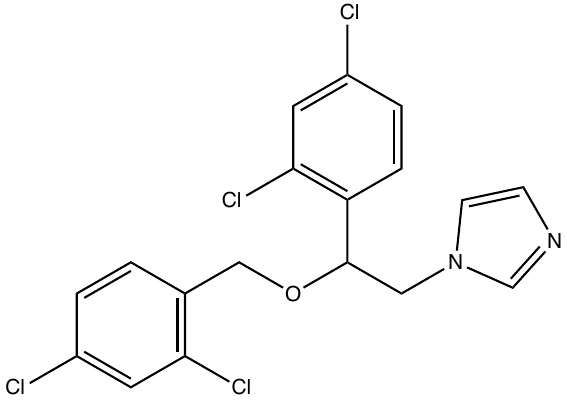 |
| Mitoxantrone | 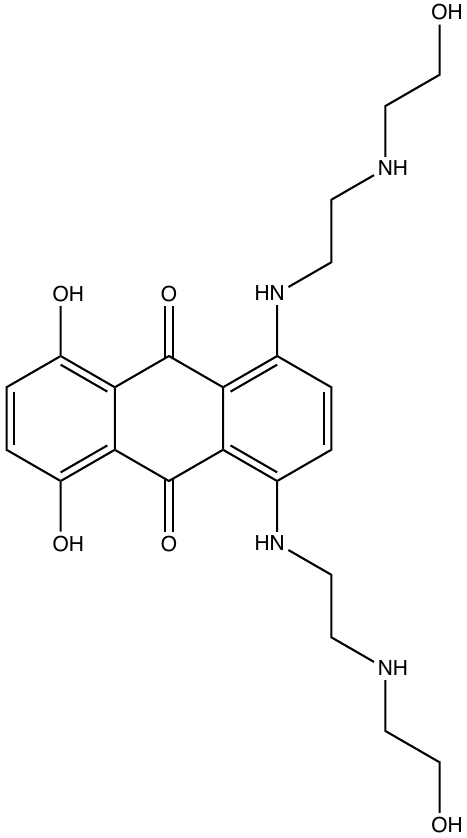 |

| **Compound Name** | **Structure** |
| --- | --- |
| Perphenazine | 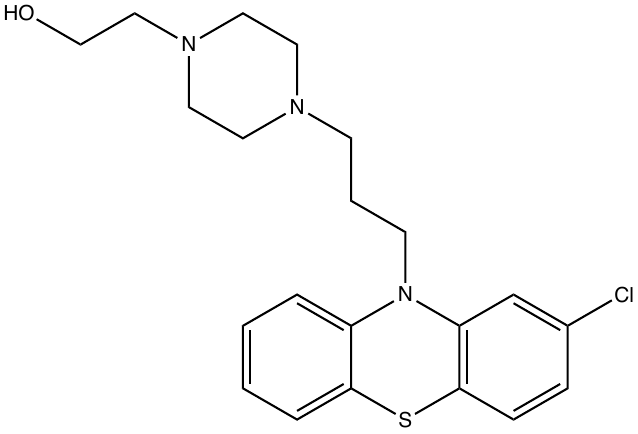 |
| Phenylmercuric acetate | 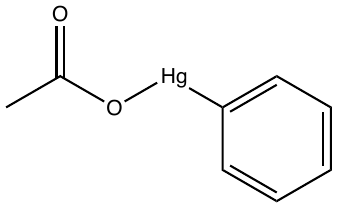 |
| Simvastatin | 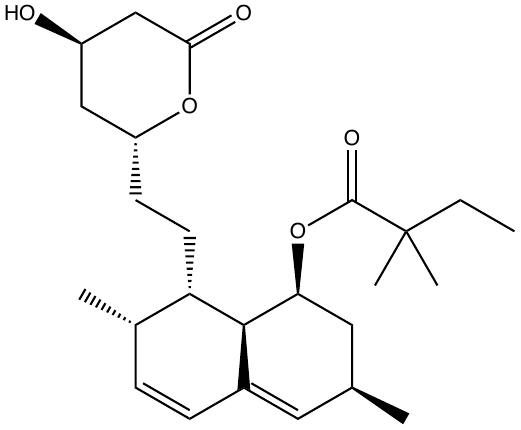 |
| Streptomycin sulfate | 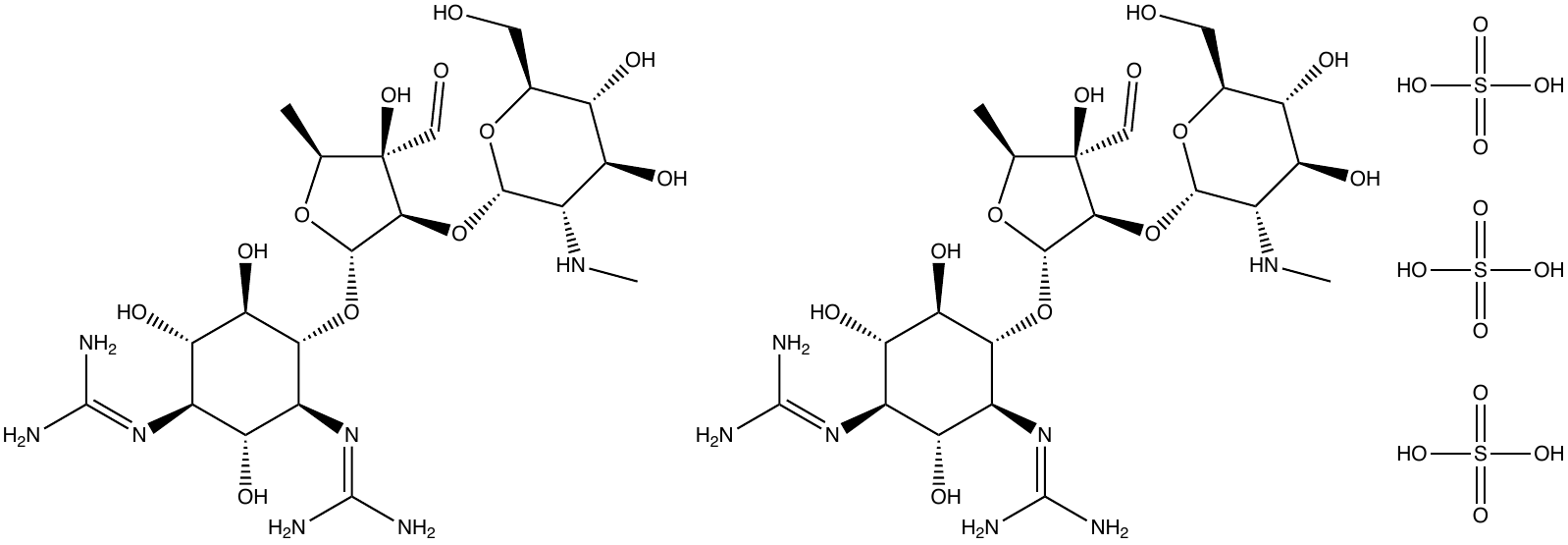 |
| Tamoxifen | 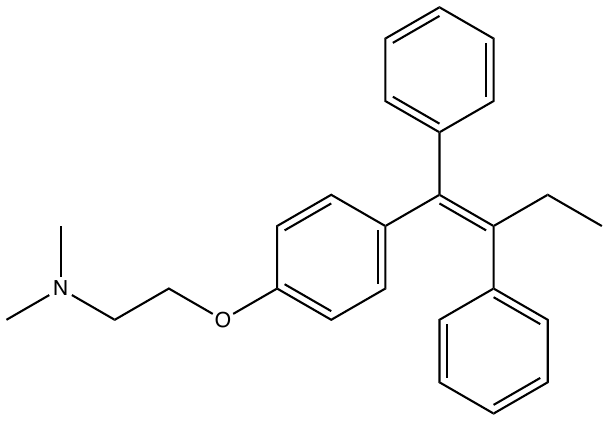 |
| Thimerosal | 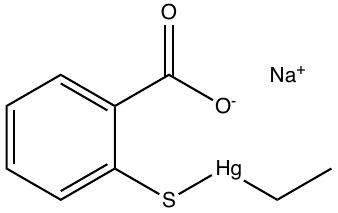 |
| Thioridazine hydrochloride | 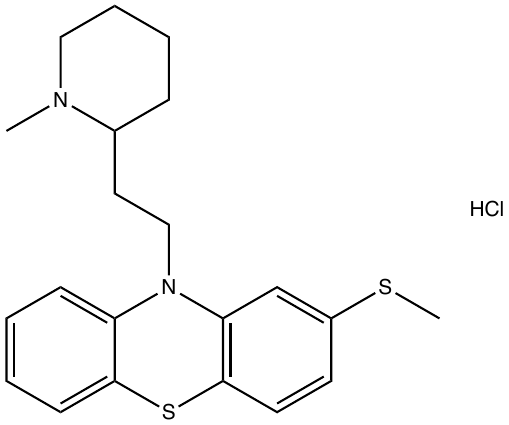 |
| Tioconazole | 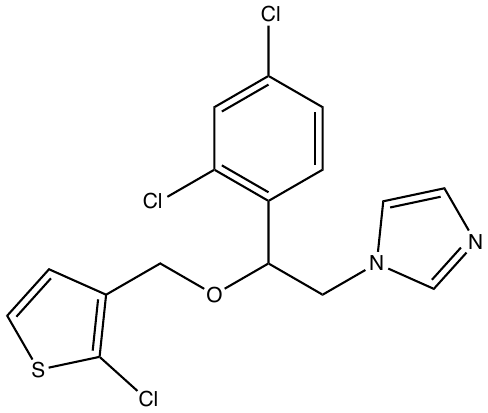 |
| Triclosan | 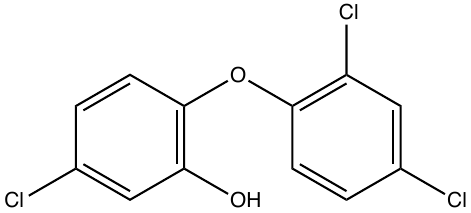 |

B)

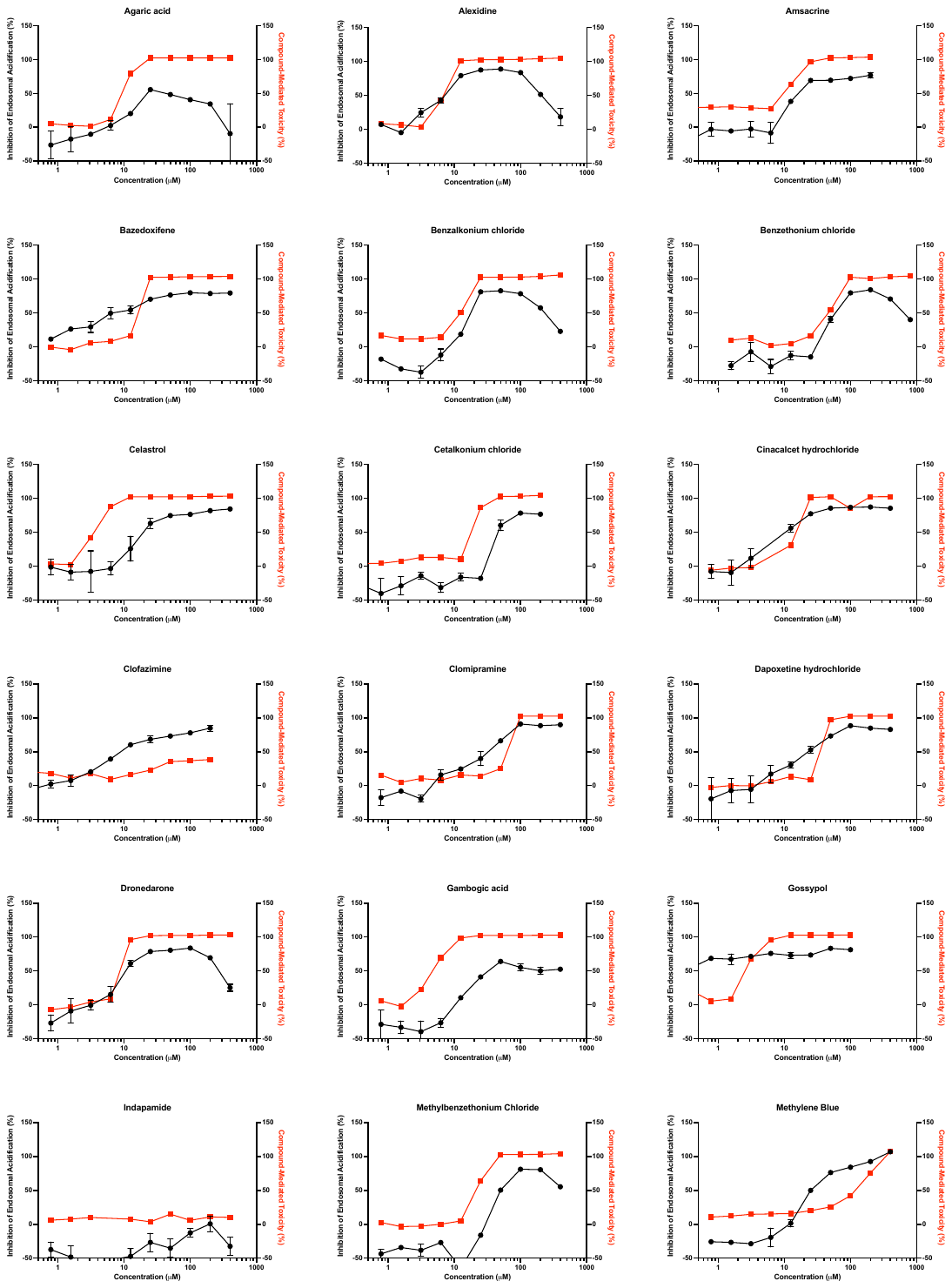

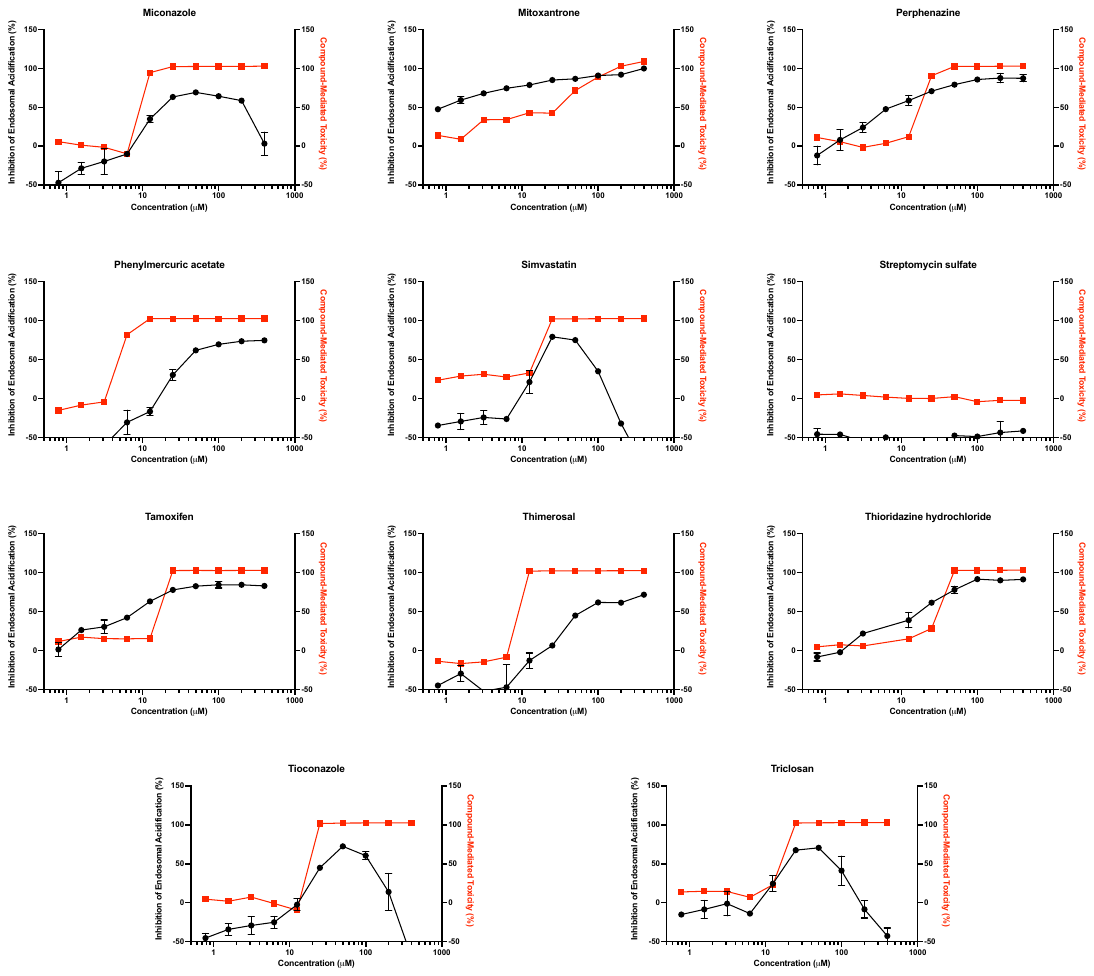

**Supplemental Table 1:** Compound screening data

| **Molecule Name** | **Percent Inhibition (%)** |
| --- | --- |
| STREPTOMYCIN SULFATE | 101 |
| INDAPAMIDE | 101 |
| DIHYDROCELASTROL | 96 |
| TRICLOSAN | 96 |
| GENTIAN VIOLET | 95 |
| ALEXIDINE HYDROCHLORIDE | 95 |
| SULOCTIDIL | 94 |
| TIOCONAZOLE | 94 |
| PERPHENAZINE | 93 |
| CELASTROL | 92 |
| THIORIDAZINE HYDROCHLORIDE | 92 |
| PERHEXILINE MALEATE | 92 |
| CINACALCET HYDROCHLORIDE | 91 |
| NONOXYNOL-9 | 91 |
| DAPOXETINE HYDROCHLORIDE | 91 |
| GAMBOGIC ACID | 90 |
| CETALKONIUM CHLORIDE | 90 |
| AGARIC ACID | 89 |
| BAZEDOXIFENE ACETATE | 89 |
| BENZALKONIUM CHLORIDE | 88 |
| AMSACRINE | 87 |
| PHENYLMERCURIC ACETATE | 86 |
| DICHLOROPHEN | 86 |
| TRIFLUPROMAZINE HYDROCHLORIDE | 86 |
| LYNESTRENOL | 86 |
| alpha-MANGOSTIN | 86 |
| METHYLENE BLUE | 85 |
| METHYLBENZETHONIUM CHLORIDE | 85 |
| DRONEDARONE HYDROCHLORIDE | 85 |
| DALBERGIONE | 85 |
| ISOPOMIFERIN | 85 |
| TOLONIUM CHLORIDE | 84 |
| HEXETIDINE | 84 |
| SIMVASTATIN | 84 |
| CLOMIPRAMINE HYDROCHLORIDE | 83 |
| TAMOXIFEN CITRATE | 82 |
| THIMEROSAL | 81 |
| MICONAZOLE NITRATE | 81 |
| THIOTHIXENE | 80 |
| GOSSYPOL | 80 |
| MITOXANTRONE HYDROCHLORIDE | 80 |
| CLOFAZIMINE | 80 |
| CETYLPYRIDINIUM CHLORIDE | 80 |
| BENZETHONIUM CHLORIDE | 80 |
| CETRIMONIUM BROMIDE | 79 |
| ISOCONAZOLE NITRATE | 79 |
| DIGITONIN | 79 |
| 4-NONYLPHENOL | 79 |
| PIPERACETAZINE | 78 |
| FLUPHENAZINE HYDROCHLORIDE | 78 |
| TRIMEPRAZINE TARTRATE | 78 |
| TRIFLUOPERAZINE HYDROCHLORIDE | 77 |
| CHLOROPHYLLIDE Cu COMPLEX Na SALT | 77 |
| AMLODIPINE BESYLATE | 76 |
| PRAZOSIN HYDROCHLORIDE | 76 |
| ROBENIDINE HYDROCHLORIDE | 76 |
| SULCONAZOLE NITRATE | 76 |
| FARNESOL | 76 |
| ECONAZOLE NITRATE | 76 |
| NICLOSAMIDE | 75 |
| PROMAZINE HYDROCHLORIDE | 75 |
| TOTAROL | 74 |
| SERTRALINE HYDROCHLORIDE | 74 |
| FENDILINE HYDROCHLORIDE | 73 |
| PROCHLORPERAZINE EDISYLATE | 73 |
| TOMATINE | 72 |
| BEPRIDIL HYDROCHLORIDE | 71 |
| PYRONARIDINE TETRAPHOSPHATE | 70 |
| ETHOPROPAZINE HYDROCHLORIDE | 70 |
| DESLORATIDINE | 70 |
| HYCANTHONE | 69 |
| AMINACRINE | 69 |
| CLOMIPHENE CITRATE | 69 |
| PRIMAQUINE PHOSPHATE | 68 |
| RAFOXANIDE | 68 |
| TOREMIPHENE CITRATE | 68 |
| FLUOXETINE | 67 |
| 18-AMINOABIETA-8,11,13-TRIENE SULFATE | 67 |
| CYPROHEPTADINE HYDROCHLORIDE | 67 |
| EBASTINE | 67 |
| BITHIONATE SODIUM | 67 |
| ACEPROMAZINE MALEATE | 67 |
| NEROLIDOL | 66 |
| TOTAROL-19-CARBOXYLIC ACID, METHYL ESTER | 66 |
| PAROXETINE HYDROCHLORIDE | 65 |
| CHLORPROMAZINE | 65 |
| CANTHARIDIN | 65 |
| BEXAROTENE | 64 |
| GUAIAZULENE | 64 |
| TERCONAZOLE | 64 |
| ARACHIDONIC ACID | 64 |
| BENAZEPRIL HYDROCHLORIDE | 64 |
| TESTOSTERONE PROPIONATE | 64 |
| MEFLOQUINE | 64 |
| METHYL DEOXYCHOLATE | 63 |
| IDEBENONE | 63 |
| VORTIOXETINE HYDROBROMIDE | 63 |
| QUINACRINE HYDROCHLORIDE | 63 |
| APREPITANT | 62 |
| CARVEDILOL PHOSPHATE | 62 |
| ACETOPHENAZINE MALEATE | 62 |
| PENFLURIDOL | 61 |
| MAPROTILINE HYDROCHLORIDE | 60 |
| AMPHOTERICIN B | 60 |
| DULOXETINE HYDROCHLORIDE | 59 |
| ABAMECTIN (avermectin B1a shown) | 59 |
| TILORONE | 59 |
| PROADIFEN HYDROCHLORIDE | 59 |
| AZELASTINE HYDROCHLORIDE | 59 |
| DROFENINE HYDROCHLORIDE | 58 |
| PYRITHIONE ZINC | 58 |
| BENZYDAMINE HYDROCHLORIDE | 58 |
| METITEPINE MALEATE | 58 |
| CHLORPROTHIXENE HYDROCHLORIDE | 57 |
| PROMETHAZINE HYDROCHLORIDE | 56 |
| CLOSANTEL | 56 |
| RALOXIFENE HYDROCHLORIDE | 56 |
| TRIMIPRAMINE MALEATE | 56 |
| CARVEDILOL | 55 |
| OXYCLOZANIDE | 55 |
| PROFLAVINE HEMISULFATE | 54 |
| CHICAGO SKY BLUE | 54 |
| PIMOZIDE | 54 |
| LOPERAMIDE HYDROCHLORIDE | 54 |
| IMIPRAMINE HYDROCHLORIDE | 54 |
| CARAZOLOL | 54 |
| DIPERODON HYDROCHLORIDE | 54 |
| METERGOLINE | 53 |
| AMITRIPTYLINE HYDROCHLORIDE | 53 |
| MEBHYDROLIN NAPHTHALENESULFONATE | 52 |
| AMIODARONE HYDROCHLORIDE | 52 |
| CYCLANDELATE | 52 |
| NEBIVOLOL HYDROCHLORIDE | 52 |
| MEPARTRICIN | 52 |
| ACRIFLAVINIUM HYDROCHLORIDE | 51 |
| HALOPERIDOL | 51 |
| FLUPIRTINE MALEATE | 50 |
| DESIPRAMINE HYDROCHLORIDE | 50 |
| BENZTROPINE MESYLATE | 50 |
| ASTEMIZOLE | 50 |
| PIZOTYLINE MALATE | 50 |
| ACRISORCIN | 50 |
| BIFONAZOLE | 50 |
| TETRANDRINE | 50 |
| BROMPERIDOL | 50 |
| TERFENADINE | 50 |
| ANDROSTERONE ACETATE | 50 |

**SUPPLEMENTAL FIGURE AND TABLE LEGENDS**

**Supplemental Figure 1. High throughput screen of the Spectrum Collection library to identify inhibitors of endosomal acidification.** (A) The calculated Z-factor from each of the 32 plates from the screen. (B) High throughput inhibition of endosomal acidification screen data. each circle represents the inhibition of a unique compound from the Spectrum Collection library tested at 40 μM. The dashed line represents the calculated hit cut off (mean plus three times the standard deviation of the data), with the red circles demonstrating inhibition values greater than the cut off. All experiment were conducted in singlet (n=1).

**Supplemental Figure 2. Dose response inhibition of endosomal acidification and compound-mediated toxicity of compounds above the hit cut off.** (A) Chemical structure of the compounds selected from the high throughput screen for further testing. (B) Comparison of the inhibition of endosomal acidification and compound-mediated toxicity by each tested compound across a dose titration range. Inhibition of endosomal acidification experiments were conducted in duplicate (n=2) and cell toxicity experiments were conducted in singlet (n=1).

**Supplemental Table 1. Inhibition of endosomal acidification by each compound from the Spectrum Collection library.**
